# Deriving connectivity from spiking activity in detailed models of large-scale cortical microcircuits

**DOI:** 10.1101/2024.06.13.598937

**Authors:** Faraz Moghbel, Muhammad Taaha Hassan, Alexandre Guet-McCreight, Heng Kang Yao, Etay Hay

## Abstract

Inferring detailed cortical microcircuit connectivity is essential for uncovering how information is processed in the brain. A common method *in vivo* uses short-lag spike cross-correlations to derive putative monosynaptic connections, but inactive neurons and correlated firing can hinder the derivation accuracy. Previous computational studies that developed methods to derive connectivity from cross-correlations employed simplified or small network models and thus did not address the above key confounds of physiological large-scale networks. We tested connectivity derivation using simulated ground-truth spiking from detailed models of human cortical microcircuits in different layers and between key neuron types. While derivation accuracy was high for cortical layer 5 microcircuits, we showed that low-firing and inactive neurons in layer 2/3 microcircuits resulted in poor performance. We then showed that general activation paradigms for layer 2/3 microcircuits led to only a moderate improvement in derivation performance, due to a trade-off between reducing the proportion of inactive neurons and increasing correlated overactive neurons. We further improved the connection derivation performance using a more refined activation paradigm leading to jittered moderate spiking, which decreased inactive neurons without incurring unwanted correlations. Our results address key physiological challenges and provide methods to improve performance in deriving connections from spiking activity in large-scale neuronal microcircuits.

## Introduction

Information processing in the brain is governed by the activity of diverse neuron types^1–3^ with specific connectivity^4–7^. Accordingly, cortical dysfunction has been associated with changes in microcircuit connectivity^8–11^. Accurately inferring the underlying cortical microcircuit connectivity is therefore essential for uncovering how information is processed in the brain in health and disease. Short-lag correlations from pairwise extracellular spiking have been commonly used to derive monosynaptic connectivity *in vivo*^12–15^, and using these methods with high-density Neuropixels probes^16,17^ can enable derivation of larger scale connectivity. However, studies have yet to address key physiological confounds of large-scale derivation such as inactive neurons^18^ and correlated network activity^19^.

There are several common methods for deriving connections, each with its advantages and limitations. Intracellular paired recordings of postsynaptic potentials (PSP) enable direct detection and measuring of synaptic connection strength^20^, but are limited in scale to at most a dozen neurons simultaneously^5^. Recent advancements allow for the switching of the neurons that are being recorded, to yield tens of neurons from a single microcircuit, but the scalability limitation remains^21^. Conversely, volume electron microscopy ^22,23^ offers a means for larger-scale connectome reconstruction of segments as large as 1 mm^3^ ^24–26^. However, structural methods are limited in their ability to quantify synaptic connection strength, relying solely on structural proxies ^27,28^. Another method is calcium imaging which measures activity from large-scale populations^29,30^, however the method’s utility is compromised by a lower temporal resolution and non-trivial relationship of the signal with spikes^31^. Short-lag spike cross-correlations between pairs of neurons from extracellular recordings can be used to derive putative monosynaptic connections^12,13,15^ and their strength^32,33^ *in vivo*, which though stemming from a functional signal would correspond to single synaptic connections between pairs of neurons. Recent high-density Neuropixels probes^16^ that record spiking in hundreds of neurons simultaneously^17,34^ offer the possibility of estimating connections at a large scale.

Physiological confounds need to be considered when deriving connectivity in realistic large-scale networks. Correlated activity of neuronal microcircuits due to oscillations, recurrent activity of neuronal populations firing together^19^ or shared inputs^35,36^, make it difficult to infer whether two neurons are directly connected or if they only appear to fire in relation to one another due common inputs^37^. These confounds can mask the evidence of true connections, leading to false negatives, or lead to spurious derived connections, thus reducing the accuracy of cross-correlation derivation methods. Inactive or silent neurons, for which spiking information is lacking, can constitute up to 30% of neurons in cortical layer 2/3 (L2/3) and ∼5% of neurons in layer 5 (L5)^18,38–40^, leading to undetectable connections. Other physiological properties such as detailed neuron morphologies, multiple synaptic contacts, and stochastic release probabilities, all contribute to a more complex relationship between synaptic inputs and spike output^41^, which may impose challenges in connection derivation from cross-correlations. Previous efforts have taken different approaches to improve neuronal connectivity derivation. Generalized linear models fit to cross-correlograms were used to improve the derivation performance and to estimate synaptic weights corresponding to PSPs^42,43^, by evaluating short-lag cross-correlations relative to a slow-wave function that fits the baseline correlations and absorbs noisy fluctuations^42,43^, instead of relative to a flat baseline as in more traditional methods^44^. These readily available tools also facilitated the automation of connection derivation at larger scale. Supervised learning algorithms have been used to infer synaptic weights from presynaptic spike trains of simulated networks^45^, and with activation paradigms to address silent neurons^46^. Other studies developed methods to derive network connectivity from multiple samples via statistical connectivity inferencing^37,47^. However, all these previous studies used simplified small network models that did not address challenges of large-scale physiological microcircuits such as silent neurons and correlated activity due to circuit dynamics.

Here we test methods for deriving putative monosynaptic connections from cross-correlations using simulated ground-truth spiking data from recent detailed large-scale models of human cortical microcircuits^48,49^. These models draw on the strengths of previous biophysically detailed models of rat and monkey microcircuits^50,51^, while having increased tractability by having fewer parameters. The models capture firing properties of key human neuron types and the synaptic connectivity between them, and reproduce network oscillatory dynamics and power spectral properties^52^, as well as a proportion of silent neurons. These models can thus serve for ground-truth simulations to assess physiological challenges on large-scale connection derivation, overcoming experimental limitations. We characterize the derivation performance for microcircuits in different cortical layers, and test general and refined activation paradigms to improve connection derivation. In addition to excitatory connections, we also examine derivation for inhibitory connections of different cell types in the cortical microcircuits.

## Results

We examined the derivation of putative monosynaptic connections from spike cross-correlations during baseline activity in cortical microcircuits with moderate vs low firing rates. We used simulated ground-truth data from our previous detailed models of human cortical L5 and L2/3 microcircuits, which have moderate and low baseline firing rates of pyramidal (Pyr) neurons, respectively (average rates over 180 minutes of baseline activity: 3.3 ± 1.0 Hz vs 0.8 ± 0.6 Hz, Cohen’s *d* = 2.96, *p* < 0.001, Figure 1A-B). We used recent tools (GLMCC toolkit^42^, see Methods) of fitting generalized linear models to cross-correlograms, featuring a slow fluctuation curve to absorb correlation trends in baseline activity over longer time courses^42,43^ (Figure 1C).

**Figure 1.**
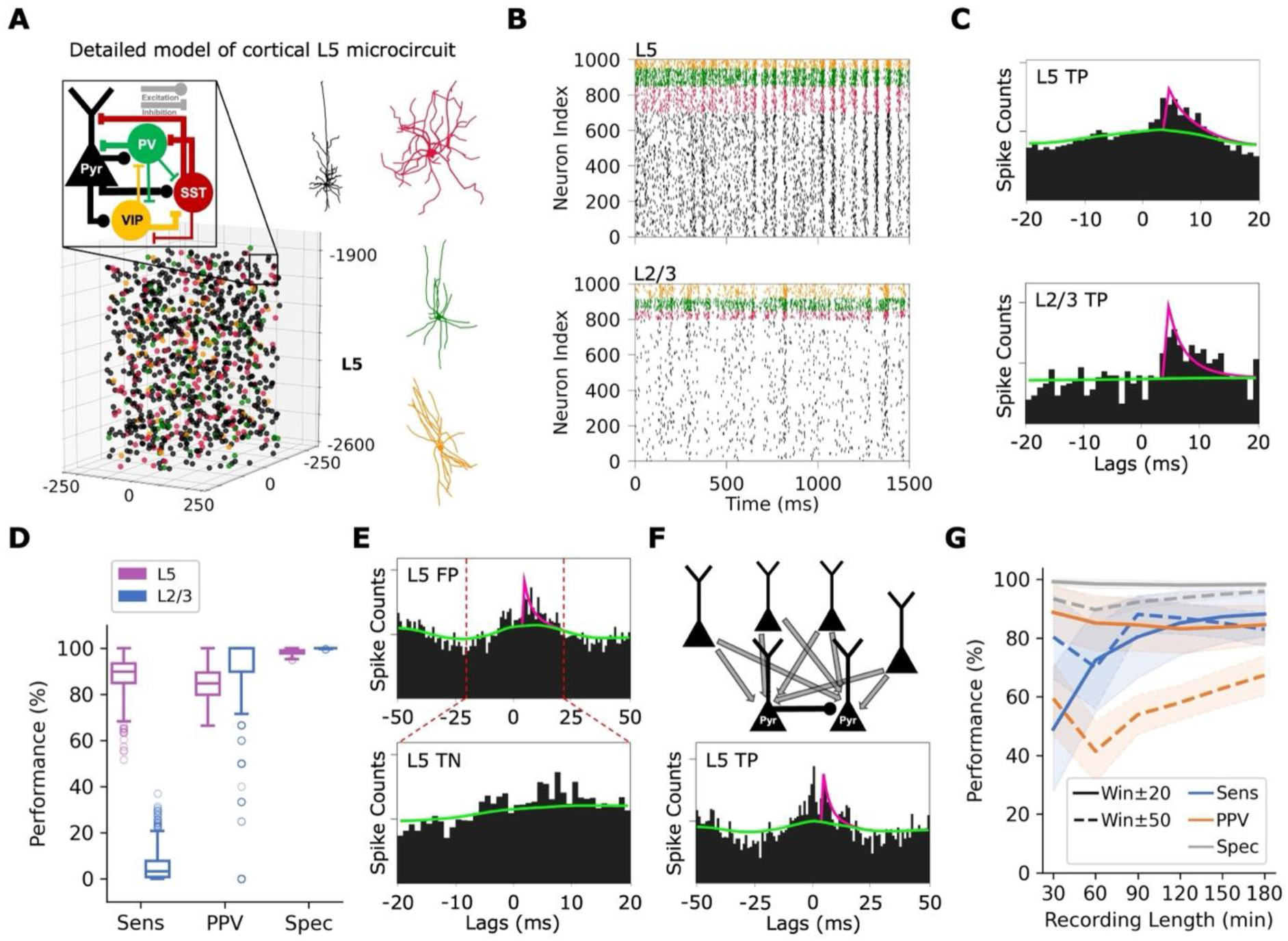
Characterizing physiological confounds when deriving excitatory connections in L5 and L2/3 microcircuits. **(A)** The detailed model of a human cortical L5 microcircuit comprised of 1,000 neurons of the key types (Pyr neurons, somatostatin-expressing (SST) interneurons, parvalbumin-expressing (PV) interneurons, and vasoactive intestinal peptide-expressing (VIP) interneurons) with detailed morphologies, distributed in a 500 x 500 x 700 μm^3^ volume. Inset: schematic diagram of the excitatory and inhibitory connections between neuron types. L2/3 microcircuits had a similar connectivity scheme, but different connection probabilities (see Table 1) and cell proportions. **(B)** Example raster plots of baseline neuronal spiking in L5 (top) and in L2/3 (bottom) used for connection derivation, color-coded according to neuron type. **(C)** Cross-correlogram, derived connection (magenta) and slow fluctuation curve (green) for an example true positive (TP) derived excitatory connection in the L5 microcircuit (top) and L2/3 microcircuit (bottom). **(D)** Derivation performance for L5 (purple, n = 700 postsynaptic Pyr neurons) and L2/3 (blue, n = 800 postsynaptic Pyr neurons) excitatory connections. Box plots show the interquartile range (IQR), median, and ±2 IQR range. The circles show outliers outside the ranges. **(E)** Example false positive (FP) L5 excitatory connection derivation at ± 50 ms window (top), which was correctly identified as a true negative (TN) using the ± 20 ms window (bottom). **(F)** Schematic of TP excitatory connection with shared inputs (top) and cross-correlogram for an example L5 Pyr connection at ± 50 ms window (bottom), which was correctly detected but also involved a peak around lag 0. **(G)** Derivation performance as a function of the length of simulated spike data in L5, using ± 50 ms (dashed lines) and ± 20 ms (solid lines) cross-correlation lag windows (lines and shaded areas show mean ± SD, respectively).

**Table 1.** Connection probabilities for L5 and L2/3 microcircuit models.

|  | **Pyr** | **SST** | **PV** | **VIP** |
| --- | --- | --- | --- | --- |
| **L5 connection probability** |  |  |  |  |
| **Pyr** | 0.09 | 0.09 | 0.09 | 0.08 |
| **SST** | 0.10 | 0.09 | 0.10 | 0.10 |
| **PV** | 0.06 | 0.16 | 0.40 | 0.10 |
| **VIP** | 0 | 0.30 | 0.20 | 0.05 |
| **L2/3 connection probability** |  |  |  |  |
| **Pyr** | 0.15 | 0.19 | 0.09 | 0.09 |
| **SST** | 0.19 | 0.04 | 0.20 | 0.06 |
| **PV** | 0.094 | 0.05 | 0.37 | 0.03 |
| **VIP** | 0 | 0.35 | 0.10 | 0.05 |

Rows correspond to the presynaptic neuron and columns correspond to the postsynaptic neuron.

Deriving excitatory connections across the moderately active L5 Pyr neurons (n = 700) had good performance accuracy (sensitivity = 88.2 ± 7.5%, positive predictive value (PPV) = 84.5 ± 6.6%, specificity = 98.3 ± 0.9%, Figure 1D). The L5 Pyr neuron baseline activity involved sufficient spiking for detecting pairwise short-lag correlations, where the slow fluctuation curve fit the long-lag dynamics well using a flatness parameter (γ) of 2 x 10^-4^ ms^-1^, and short-lag peaks were fit well when testing for connections in a 4 – 8 ms window (Figure 1C). In contrast to the moderately active L5, the lack of spikes in low-firing L2/3 Pyr neurons made it more difficult to detect short-lag peaks compared to baseline activity (Figure 1C), resulting in poor connection derivation performance (sensitivity = 5.4 ± 6.2%, n = 800 postsynaptic neurons, Figure 1D).

Compared to the above parameters for L5, using an earlier synaptic delay, such as a traditional d_s_ = 2 ms^12,13,15^ led to poorer PPV (36.5 ± 7.4, Cohen’s *d* =-6.82, *p* < 0.001) and lower sensitivity (79.3 ± 7.1, Cohen’s *d* =-0.96, *p* < 0.001). Using a flatter slow fluctuation curve e.g. γ = 1 x 10^-4^ ms^-1^ led to increased spurious derivations (PPV = 61.4 ± 16.4, Cohen’s *d* =-1.85, *p* < 0.001), while a more oscillatory curve e.g. γ = 6 x 10^-4^ ms^-1^ reduced sensitivity (29.9 ± 7.8, Cohen’s *d* =-6.88, *p* < 0.001). We tested these parameters on another L5 microcircuit model, with a different random instantiation of the connectivity and background inputs. Derivation performance was similar in both microcircuits at 30 min of baseline activity (sensitivity = 49.0 ± 21.2% vs 50.3 ± 21.3, Cohen’s *d* = 0.06, *p* = 0.29; PPV = 88.8 ± 8.8 vs 87.0 ± 9.6, Cohen’s *d* =-0.19, *p* < 0.001), thus supporting reproducibility of the results.

While testing for connections in the 4 – 8 ms lag window, we used an overall analysis window of ± 20 ms cross-correlation lags to inform the slow wave function with baseline correlation values, as the L5 microcircuit model activity oscillated in the beta frequency range in line with previous experiments^53^. The presence of multiple oscillation peaks at longer cross-correlogram analysis windows such as ± 50 ms or ± 200 ms, traditionally used in previous works^12,19,42^, skewed the tool’s estimation of the baseline correlation with the slow wave function and led to worse sensitivity and PPV (sensitivity = 83.0 ± 5.7%, Cohen’s *d* =-0.79, *p* < 0.001; PPV: 67.3 ± 6.8%, Cohen’s *d* =-2.57, *p* < 0.001, Figure 1E,G). Using the analysis window of ± 20 ms, shorter than the oscillation period of ∼50 ms corresponding to the peak frequency of 17.0 Hz in the power spectral density, improved connection derivation by ensuring that the oscillations were not factored into the baseline estimation and overall analysis. While the slow fluctuation curve helped derivation, it could not sufficiently mitigate the effects of highly correlated activity, seen especially in the longer windows (Figure 1E), regardless of the parameter values. For example, even using a wave period that closely fit the long-lag dynamics in the analysis window (see performance for γ = 6 x 10^-4^ ms^-1^ above) did not improve the derivation, as it increased the proportion of false negatives. Excluding connections that peaked around lag 0 (between-1 and 1 ms, indicating correlated activity rather than necessarily a connection) did not improve derivation either, as it improved PPV (80.2 ± 7.5%, Cohen’s *d* = 1.80, *p* < 0.001) at the cost of reduced sensitivity (59.4 ± 13.1%, Cohen’s *d* =-2.34, *p* < 0.001). The peaks around lag 0, indicative of shared rhythmicity, were thus also present in many true connections, and thus ruling out connections based on a peak at lag 0 did not improve connection derivation (Figure 1F).

To determine the robustness of our results and sufficiency of the spike data length, we characterized performance as a function of data length. Connection derivation performance using shorter data (30 or 60 minutes) was not satisfactory, thus requiring longer spike data. There was a steady improvement in derivation performance with the length of simulated data from 30 minutes up to 120 minutes (sensitivity = 85.2 ± 10.9%, Cohen’s *d* = 2.15, *p* < 0.001, Figure 1G), beyond which the improvement was only marginal (120 min vs 180 min: sensitivity = 85.2 ± 10.9% vs 88.2 ± 7.5%, Cohen’s *d* = 0.32, *p* < 0.001, Figure 1G). Furthermore, the distribution of indegree for derived Pyr→Pyr connections was similar to that of the true connections (median = 67.0, IQR = 17.0 vs median = 63.0, IQR = 10.0, Cohen’s *d* = 0.33), although it had a wider range with a slightly heavier right tail (95% range: [41.0, 91.5] vs [48.0, 77.5]).

We therefore tested a general activation of L2/3 Pyr neurons by increasing their background input (hereafter referred to as L2/3 general-Pyr), thus increasing the mean firing rate to 3.3 ± 2.4 Hz, close to that in L5 (Figure 2A). Derivation performance for the L2/3 general-Pyr paradigm had higher sensitivity (50.9 ± 14.0%, Cohen’s *d* = 3.91, *p* < 0.001, n = 800 neurons, Figure 2B), although still low compared to the derivation from the L5 microcircuit. While the general-Pyr activation decreased PPV to some extent (L2/3 baseline: PPV = 88.7 ± 24.0%, n = 800 neurons; L2/3 general-Pyr: PPV = 70.4 ± 14.2%, Cohen’s *d* = - 0.93, *p* < 0.001, n = 800 neurons, Figure 2B), the increase in sensitivity was much larger, therefore the derivation performance was overall better and more balanced than at baseline.

**Figure 2.**
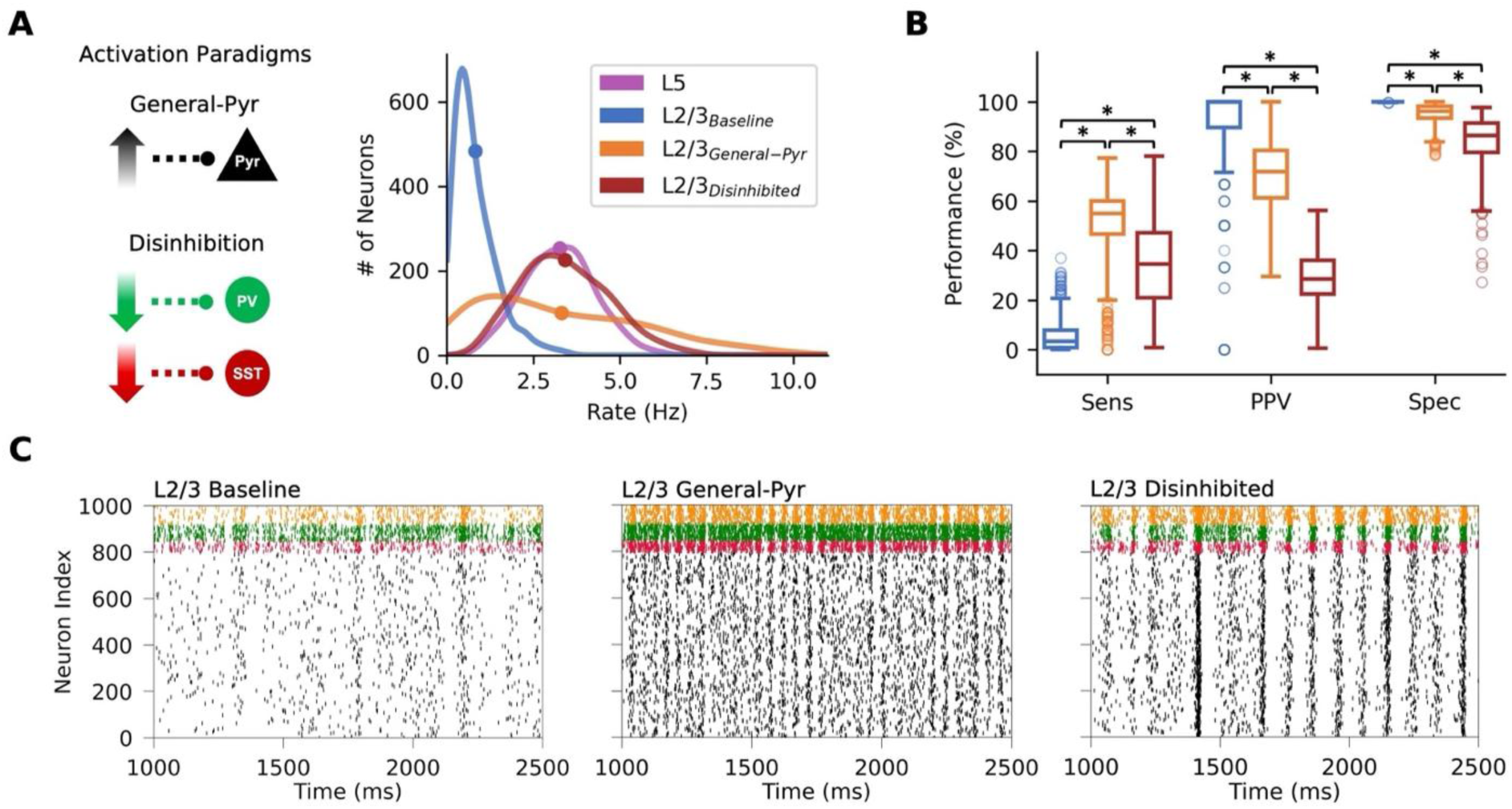
Moderate improvement in derivation of L2/3 connections by activation of Pyr neurons. **(A)** Left: The paradigms for activation of Pyr neurons in L2/3 microcircuits, either generally activating Pyr neurons or by disinhibition. Right: Distribution of average Pyr neuron spike rates in L5 baseline (purple, n = 700), L2/3 baseline (blue, n = 800), L2/3 general-Pyr (orange, n = 800), and L2/3 disinhibited (red, n = 800). Curves were smoothed using a Gaussian kernel. Circles denote the mean. **(B)** Derivation performance for excitatory connections in L2/3 microcircuits at baseline, and in the general-Pyr and disinhibited paradigms (color-coded as in A). Box plots show the IQR, median, and ±2 IQR range. The circles show outliers outside the range. Asterisk denotes *p* < 0.001 and Cohen’s *d* > 0.5. **(C)** Example raster plot of simulated spiking in L2/3 microcircuits at baseline (left), and in the general-Pyr (middle) and disinhibited (right) paradigms, color-coded according to neuron type.

A likely reason for the moderate improvement in L2/3 general-Pyr was indicated by the distribution of average Pyr neuron spike rates (Figure 2A). While the Pyr neurons in L2/3 general-Pyr had much less inactive neurons compared to baseline L2/3, there were still 18% (n = 147) inactive neurons firing at < 1 Hz, and the sensitivity in deriving connections for these neurons was poorer than average (10.1 ± 10.0%, Cohen’s *d* =-2.80, *p* < 0.001). There was also a larger right tail compared to L5, with a considerable proportion (25%, n = 197) of overactive neurons firing at > 5 Hz, for which the sensitivity in connection derivation was poorer than average too (13.8 ± 7.9%, Cohen’s *d* =-2.63, *p* < 0.001). The main reason for the low sensitivity in the case of overactive neurons was a large proportion of false negatives that were estimated as non-connections due to peaks at lag 0 in the cross-correlograms, resulting from higher spike correlations between the overactive neurons than average (0.008 ± 0.008 vs 0.005 ± 0.008, Cohen’s *d* = 0.4, *p* < 0.001). A reason for the lower PPV was that the increased firing of the overactive neurons involved increased recurrent oscillatory activity, characterized by increased peak frequency (from 10.3 Hz to 20.8 Hz) and increased power (from 1e^-5^ ± 8e^-6^ spikes^2^/Hz to 5e^-4^ ± 3e^-4^ spikes^2^/Hz, Cohen’s *d* = 2.46, *p* < 0.001) of Pyr neuron spiking, which led to short-lag peaks in the cross-correlograms irrespective of connectivity and thus spurious connection derivation. In comparison, the remaining 57% (n = 456) of the neurons, which fired between 1 – 5 Hz, showed better connection derivation compared to average performance (sensitivity 69.9 ± 9.0%, Cohen’s *d* = 1.43, *p* < 0.001).

We therefore tested another general activation paradigm that yielded a distribution of average L2/3 Pyr neuron spiking similar to L5 in addition to a similar mean Pyr firing rate of 3.4 ± 1.3 Hz. This paradigm involved a disinhibition of Pyr neurons by decreasing background input to PV and SST interneurons, together with a small decrease in Pyr background inputs. However, this activation by disinhibition led to more rhythmic activity of Pyr neuron spiking with multiple peak frequencies in the power spectral density (from 10.3 Hz to both 9.3 Hz and 18.8 Hz) and increased peak power (from 1e^-5^ ± 8e^-6^ spikes^2^/Hz to 4e^-3^ ± 2e^-3^ spikes^2^/Hz, Cohen’s *d* = 3.25, *p* < 0.001, and 2e^-3^ ± 1e^-3^ spikes^2^/Hz, Cohen’s *d* = 2.77, *p* < 0.001), and more correlated firing (0.06 ± 0.02 vs 0.0008 ± 0.007, Cohen’s *d* = 3.65, *p* < 0.001, Figure 2C), which resulted in increased spurious (false positive) derivation and thus poor PPV compared to baseline (29.1 ± 9.8%, Cohen’s *d* =-3.25, *p* < 0.001, n = 800 neurons, Figure 2B). The disinhibition led to a smaller increase in sensitivity from baseline (34.3 ± 17.3%, Cohen’s *d* = 2.22, *p* < 0.001, n = 800 neurons, Figure 2B), compared to the L2/3 general-Pyr paradigm (Cohen’s *d* =-1.02, *p* < 0.05) and thus was less successful.

We then tested a more refined activation paradigm, targeting a subset of 100 Pyr neurons and inducing moderate and jittered spiking (Figure 3A). This led to an increased mean firing rate (3.0 ± 1.1 Hz) only in the subset population and had little effect on the activity of the non-activated population (Figure 3B). The subset activation paradigm yielded a firing rate distribution similar to L5, with less inactive and overactive neuron populations compared to the same subset in the L2/3 general-Pyr paradigm (inactive portion = 3% vs 14%, overactive portion = 6% vs 19%, respectively, Figure 3C). Consequently, the subset activation paradigm improved the sensitivity of derivation performance compared to L2/3 general-Pyr (74.0 ± 15.2%, Cohen’s *d* = 1.06, *p* < 0.001, Figure 3D) while maintaining a similarly fair PPV performance (73.4 ± 16.0%). A stronger subset activation (e.g. leading to 4.8 Hz) was less optimal due to poorer PPV (28.7 ± 8.3%). The distribution of indegree for derived Pyr→Pyr connections of the activated subset was similar to that of the true connections (median = 15.0, IQR = 7.0, 95% range = [5.5, 27.1] vs median = 14.5, IQR = 5.0, 95% range = [8.5, 21.0], Cohen’s *d* = 0.186, *p* = 0.2). We tested the stimulation protocol on another L2/3 microcircuit model, with a different random instantiation of the connectivity and background inputs, comparing derivation performance at 30 min of activity. Performance was similar in both microcircuits (sensitivity = 35.7 ± 16.5 vs 37.3 ± 14.7, Cohen’s *d* = 0.11, *p* = 0.46; PPV = 76.9 ± 18.1 vs 79.7 ± 16.7, Cohen’s *d* = 0.17, *p* = 0.29), thus supporting reproducibility of the results.

**Figure 3.**
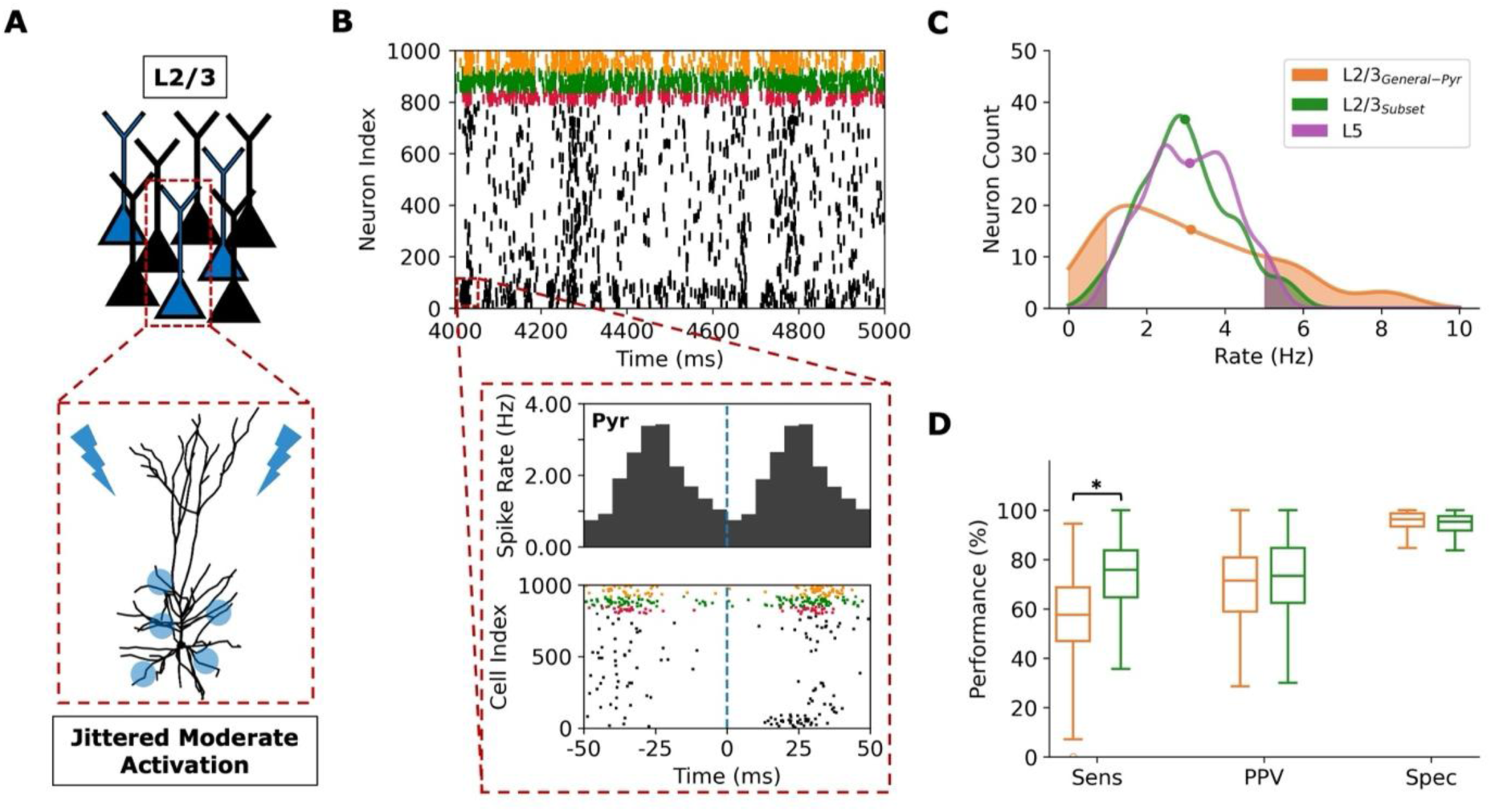
Improved true positive derivation of L2/3 connections by jittered moderate activation of Pyr neuron subset. **(A)** Schematic of simulation paradigm, where a subset of L2/3 Pyr neurons (blue) were activated repetitively by jittered moderate inputs (20 ms jitter, 50 ms cycles for 180 minutes). Inset: Schematic of synaptic contacts of the input along the basal dendrites. **(B)** Example raster plot of spiking in the L2/3 microcircuit during two cycles of the activated subset (first 100 neurons). Inset: peristimulus time histogram (top, n = 200 activation cycles) and example raster plot of spike response (bottom). A dashed line denotes stimulus onset. **(C)** Distribution of average Pyr neuron spike rates of the 100 L2/3 Pyr neuron subset (green) compared to in the L2/3 general-Pyr paradigm (orange) and a subset of 100 L5 Pyr neurons (black). The shaded regions represent the proportion of inactive (< 1 Hz) and overactive (> 5 Hz) neurons in each paradigm. Curves were smoothed using a Gaussian kernel. Circles denote the mean. **(D)** Derivation performance for excitatory connections for the 100 L2/3 Pyr neuron subset vs the general-Pyr paradigm (color-coded as in C). Box plots show the IQR, medians and ±2 IQR. Asterisk denotes *p* < 0.001 and Cohen’s *d* > 0.5.

We next compared derivation for connections from different inhibitory interneuron types onto Pyr neurons. PV→Pyr connections targeted proximal regions of the Pyr neuron (basal dendrites and soma) and could be derived using a short timescale like the Pyr→Pyr connections (τ = 4 ms in both L5 and L2/3, Figure 4A). Since SST interneurons targeted distal regions (apical dendrites), deriving their connections required using a longer decay time constant (τ = 8 ms and τ = 15 ms for SST→Pyr in L5 and L2/3, respectively, Figure 4B). For the L5 microcircuit, inhibitory connections were derived with high accuracy for both PV→Pyr (sensitivity = 100.0 ± 0.0%; PPV = 99.4 ± 1.2%; n = 100 presynaptic neurons, Figure 4C) and SST→Pyr (sensitivity = 92.5 ± 3.9%; PPV = 94.7 ± 4.8%; n = 150 presynaptic neurons, Figure 4D). In the L2/3 microcircuit at baseline, however, derivation performance was lower for both PV→Pyr connections (sensitivity = 52.9 ± 8.3%; PPV = 80.4 ± 5.6%; n = 70 presynaptic neurons, Figure 4C) and SST→Pyr connections (sensitivity = 69.1 ± 6.5%; PPV = 97.8 ± 1.8%; n = 50 presynaptic neurons, Figure 4D), although better than for Pyr→Pyr connections.

**Figure 4.**
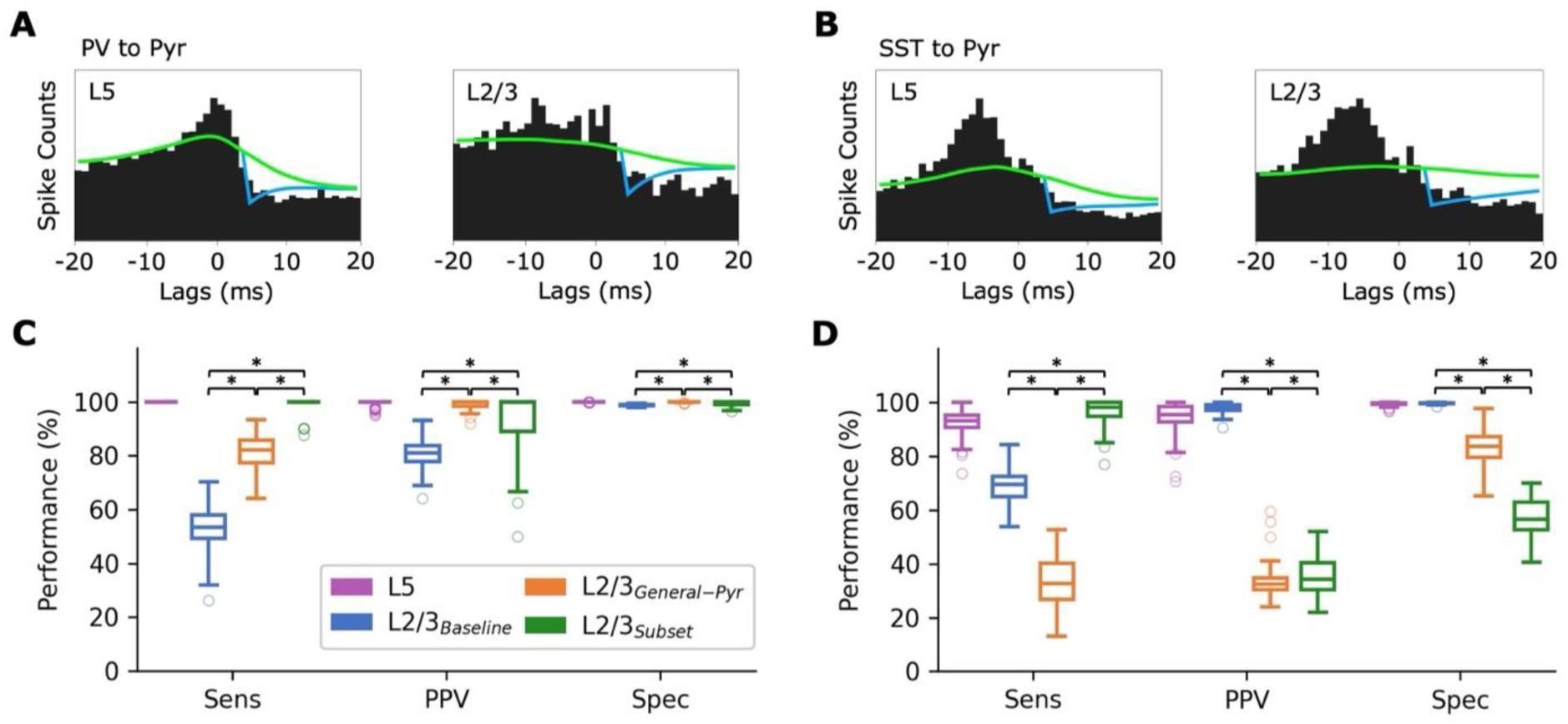
Connection derivation for PV vs SST inhibitory connections. **(A)** Cross-correlogram, derived connection (cyan) and slow fluctuation curve (green) for example derivations of true positive PV→Pyr connections in L5 (left) and L2/3 (right) microcircuits (τ = 4 ms for both). **(B)** Same as A, but for SST→Pyr connections (τ = 8 ms and τ = 15 ms in L5 and L2/3, respectively). **(C)** Derivation performance for PV→Pyr inhibitory connections in the different paradigms. **(D)** Same as C, but for SST→Pyr connections. Box plots show IQR, medians and ±2 IQR range. The circles show outliers outside the ranges. Asterisk denotes *p* < 0.001 and Cohen’s *d* > 0.5.

Derivation performance for these connections was averaged across presynaptic interneurons instead of postsynaptic pyramidal neurons to ensure adequate sample sizes. The low sensitivity in L2/3 at baseline was likely mostly due to the low firing rates in Pyr neurons, as SST and PV interneurons fired at moderate rates (5.7 ± 0.9 Hz and 10.2 ± 2.9 Hz, respectively). The higher accuracy in SST→Pyr derivation compared to PV→Pyr in L2/3 baseline was likely due to the use of a longer decay time constant and thus larger information about the effects on postsynaptic neuron spiking in cross-correlations. Using the shorter τ = 4 ms for L5 SST→Pyr connections resulted in poor sensitivity (43.4 ± 10.6%, Cohen’s *d* =-6.11, *p* < 0.001), while using a longer τ = 15 ms resulted in more spurious derivation (PPV = 66.2 ± 8.6%, Cohen’s *d* =-4.07, *p* < 0.001). In L2/3, SST→Pyr connection derivation using τ = 4 or 8 ms resulted in poor sensitivity (2.9 ± 1.4%, Cohen’s *d* =-13.99, *p* < 0.001; 26.5 ± 5.7%, Cohen’s *d* =-6.89, *p* < 0.001, respectively), requiring the longer τ = 15 ms. For PV→Pyr connections, the longer τ = 15 ms was not beneficial, resulting in poorer PPV (67.5 ± 4.6%, Cohen’s *d* =-2.50, *p* < 0.001).

The L2/3 general-Pyr activation improved derivation performance in PV→Pyr connections (sensitivity = 80.9 ± 7.0%, Cohen’s *d* = 3.62, *p* < 0.001; PPV = 99.1 ± 1.6%, Cohen’s *d* = 4.50, *p* < 0.001, Figure 4C), due to increased postsynaptic Pyr firing rates. The sensitivity was still lower than in L5 derivation, likely due to issues with the inactive and overactive postsynaptic Pyr neurons described above for Pyr→Pyr connection derivation. Derivation performance for SST→Pyr connections decreased considerably in the L2/3 general-Pyr paradigm instead of improving (sensitivity = 33.2 ± 9.6%, Cohen’s *d* =-4.35, *p* < 0.001; PPV = 33.9 ± 6.5%, Cohen’s *d* =-13.33, *p* < 0.001, Figure 4D). This was due to a large increase in SST firing rates in the L2/3 general-Pyr paradigm (24.5 ± 5.4 Hz), accompanied by increased peak frequency (from 12.3 Hz to 20.8 Hz) and power (from 3e^-6^ ± 2e^-6^ spikes^2^/Hz to 1e^-4^ ± 7e^-5^ spikes^2^/Hz, Cohen’s *d* = 2.50, *p* < 0.001) of SST interneurons spiking and higher correlated firing of SST and Pyr neurons (SST-Pyr: 0.02 ± 0.009 vs 0.002 ± 0.007, Cohen’s *d* = 1.65, *p* < 0.001; SST-SST: 0.06 ± 0.009 vs 0.009 ± 0.007, Cohen’s *d* = 5.48, *p* < 0.001), leading to spurious connection derivation and missed connections masked by correlations due to shared rhythmicity. The disinhibition paradigm resulted in even poorer derivation performance for both types of connections (PV→Pyr: sensitivity = 1.3 ± 1.3%, Cohen’s *d* =-8.62, *p* < 0.001; SST→Pyr: sensitivity = 1.6 ± 1.0%, Cohen’s *d* =-14.41, *p* < 0.001), due to even higher correlated firing in the microcircuit (SST-SST: 0.2 ± 0.01, Cohen’s *d* = 19.43, *p* < 0.001; PV-PV: 0.2 ± 0.02 vs 0.01 ± 0.008, Cohen’s *d* = 13.23, *p* < 0.001; SST-Pyr: 0.09 ± 0.02, Cohen’s *d* = 4.96, *p* < 0.001; PV-Pyr: 0.1 ± 0.03 vs 0.003 ± 0.007, Cohen’s *d* = 4.70, *p* < 0.001). The disinhibition also led to an increase in rhythmicity, characterized by multiple peaks in the spiking power spectral density of SST interneurons (from 3e^-6^ ± 2e^-6^ spikes^2^/Hz to 3e^-4^ ± 1e^-4^ spikes^2^/Hz, Cohen’s *d* = 3.25, *p* < 0.001, and 1e^-4^ ± 7e^-5^ spikes^2^/Hz, Cohen’s *d* = 2.64, *p* < 0.001) and PV interneurons (from 7e^-6^ ± 4e^-6^ spikes^2^/Hz to 5e^-4^ ± 2e^-4^ spikes^2^/Hz, Cohen’s *d* = 3.14, *p* < 0.001, and 3e^-4^ ± 1e^-4^ spikes^2^/Hz, Cohen’s *d* = 2.67, *p* < 0.001).

Compared to L2/3 baseline and L2/3 general-Pyr, the subset activation paradigm led to further improvement of PV→Pyr connection derivation for the activated subset of Pyr neurons, yielding performance similar to L5 (sensitivity = 99.4 ± 2.5%, Cohen’s *d* = 3.58, *p* < 0.001; PPV = 92.6 ± 10.0%, Cohen’s *d* = 1.00, *p* < 0.001, Figure 4C). On the other hand, for SST→Pyr connections, whereas the subset activation similarly improved sensitivity (96.2 ± 5.1%, Cohen’s *d* = 2.74, *p* < 0.001, Figure 4D), PPV was poor as in general-Pyr activation (35.0 ± 6.7%, Cohen’s *d* =-11.83, *p* < 0.001). Therefore, similarly to excitatory connections, the subset activation paradigm yielded better derivation performance for PV→Pyr and SST→Pyr connections compared to the general activation.

## Discussion

In this work, we examined physiological challenges of deriving putative monosynaptic neuronal connectivity from short-lag cross-correlations of spiking activity, using detailed models of large-scale human cortical microcircuits in different layers. We showed that silent and low-firing neurons in L2/3 led to poor derivation performance, and a general activation of the cortical microcircuit to increase firing rates yielded only a moderate improvement in connection derivation due to a trade-off between reducing inactive neurons and increasing correlated overactive neurons. We improved derivation using a more refined activation paradigm that induced moderate and jittered spiking with a reduced proportion of inactive and overactive neurons. In addition, we found that SST interneuron connections required a longer range of cross-correlation lags compared to PV interneurons, due to their distal synaptic contacts. Deriving connectivity with spike cross-correlations is currently the only method that can be applied at large-scale in the living brain, and it also enables linking microcircuit connectivity with properties of the recorded large-scale spiking dynamics in different brain states and conditions. Our results provide experimentally testable insights that can reduce the challenge of undetectable silent neurons by activation, and the challenge of spurious derivation by using jittered and moderate activation, to improve derivation of putative monosynaptic neuronal connections from spiking activity recorded by dense multi-electrodes *in vivo* and at large scale.

The general microcircuit activation paradigms in the L2/3 microcircuit aimed to increase the average firing rate of Pyr neurons to resemble L5, which was active enough to yield good derivation. These activations could correspond to a broad modulation of network activity e.g. by a pharmacological or external simulation^54^. However, unlike the normal distribution of moderate firing rates in L5, the Pyr neurons in the L2/3 general-Pyr activation paradigm had populations of overactive neurons and remaining underactive neurons hindering derivation performance. General activation of Pyr cells via disinhibition led to highly oscillatory and correlated pyramidal activity, in line with previous studies of reduced PV interneuron inhibition^55^, whereby inhibition primarily occurred as feedback following Pyr neuron firing. Overall, the reduced connection derivation performance due to correlated and rhythmic activity suggests a limited use of general activation paradigms, as increased recurrent activity of neuronal populations involves cross-correlation peaks near lag 0 which mask the effect of connections, and also introduces short-lag peaks leading to spurious derivations. The improvement we achieved using the more refined subset activation protocol for jittered moderate firing indicates the benefit of diverse experimental paradigms e.g. presentations of diverse visual stimuli^34^ or somatosensory^49,56^ or motor tasks^57,58^, to activate subsets of neurons moderately and variably to further improve connection derivation performance. Due to the required recording length of more than an hour with a moderate firing rate of 3 – 5 Hz for good derivation, in line with previous studies^42^, future studies can investigate the effect of plasticity on the derivation, although it should primarily affect the weight of the connections, and could be mitigated using diverse stimulus presentations instead of repetitive tasks. The subset activation was also able to improve derivation of PV→Pyr connections, but was less effective for deriving SST→Pyr connections, likely because the activation involved basal stimulation of Pyr neurons, for which PV interneuron inhibition is more relevant.

When deriving monosynaptic connections from short-lag correlations, oscillatory firing, neuronal ensembles and phasic activation (e.g. PV and SST interneuron recruited by Pyr neuron firing) are all confounds rather than features, since they mask and add noise to the short-lag signatures of the connections. Overcoming the challenge of microcircuit oscillations required shortening the cross-correlogram analysis window to less than the oscillation period, as it enabled the slow wave of the tool to better fit the baseline fluctuations near lag 0 and thus better detect connections on top of the baseline cross-correlation. Trying to improve PPV by excluding connections with peaks near lag 0 had a high cost on sensitivity, indicating that while correlated activity in cross-correlograms could lead to spurious derivations, a sizeable proportion of neuron pairs that fired together were also connected. Thus, the derivation would not benefit from excluding them, presenting a trade-off between sensitivity and PPV in large-scale neuronal networks with populations that fire together. Although we did not model shared inputs when activating or for background input, there were shared inputs within the microcircuits that partly contributed to the poor derivation when the neurons were firing together. The moderate and jittered L2/3 subset activation performed better because it overcame the common input problem that is exacerbated by more correlated and rhythmic activity. Future studies can model shared activation and background inputs between neurons^36,37,59^ to refine the assessment of activation paradigms for connection derivation. Furthermore, the tool’s components were not sufficient to fit the overall shape of the cross-correlogram trough of inhibitory SST→Pyr connections which involved a slower rise time, resulting in moderate derivation performance and highlighting the tool’s limitation of having only a single time constant parameter. Future studies should try to improve the tool by including both rise and decay time constants in the coupling filter.

Connection derivation in our microcircuit models required a slightly longer synaptic delay compared to previous studies^13,42^. A possible reason is that traditional parameters were based on rodent data, where neurons have shorter and less complex dendritic arbors than human neurons^60–62^, and thus the delay between presynaptic spike and PSP at soma is shorter. While the cross-correlogram lag range used for connection derivation was comparable to previous studies, good derivation of L2/3 SST→Pyr connections necessitated a longer lag range compared to PV→Pyr connections due to the attenuation of SST IPSP from the distal apical synapses to the soma^63,64^. Deriving L5 SST→Pyr connections also required a long range but to a lesser extent, since human L5 Pyr neuron apical dendrites were less branched than those of L2/3 Pyr neurons in the reconstructed morphologies we used, and also on average as reported previously^48,49^. The proximal locations of PV→Pyr synapses resulted in stronger connections compared to SST→Pyr as seen experimentally^65,66^, explaining the overall better derivation performance in L5, and a better improvement when activating L2/3. In this work we focused on connection detection rather than weight derivation, since the microcircuit models had the same conductance for all connections of a given type, with only the contact locations varying, resulting in a low range of synaptic weights. Future models should account for wider synaptic weight distributions as seen in the cortex^4^ to study derivation performance in terms of connection weights. Improving derivation of large-scale connectivity and weight estimation for inhibitory connections will enable a better characterization and modeling of their role in health and prediction of their contribution in disease. Distal dendritic-targeting SST→Pyr connections modulate the processing of top-down (cortico-cortical) inputs, maintain baseline activity and signal-to-noise ratio via lateral inhibition^48^, and mediate slower network rhythms^67,68^. Reduced SST interneuron connection strength is implicated as a mechanism of major depressive disorder^8,9^. Proximal dendritic-targeting PV→Pyr connections modulate faster network rhythms^69,70^ and their dysfunction is implicated primarily in schizophrenia^10,11^.

While constraints for the L5 and L2/3 microcircuit models came mostly from human data, some constraints including connection probabilities and synaptic amplitudes were partly based on rodent data^48,49^, but the models captured prototypical cortical microcircuit properties that are preserved across species such as intrinsic firing properties and connectivity patterns of cell types^71^. Thus, we expect our results and insights to hold regardless of the particular species.

Future studies can further integrate human data constraints as they become available^6,26,72^, to enhance species specificity.

Future studies can refine our estimates of derivation performance by addressing issues that would arise from extracellular recordings such as noise^19,73,74^ and spike sorting^75–77^. In addition, including a larger proportion of silent neurons as suggested experimentally^18,38–40^ and shared inputs among the neurons in the microcircuits^35,36^ will help refine the estimated performance and the necessary activation paradigms. Future studies can refine the assessment of activation paradigms in multi-layer microcircuit models that include L2/3, L4, and L5, to assess derivation of interlaminar connections (e.g. L4 inputs onto L2/3 and connections between L5 and L2/3^29^), although we expect our general and subset activation insights to mostly hold. Future studies can also probe the effects of morphology^41^ or physiological synaptic features such as multiple contacts, failure probability^50,78^ and short-term dynamics on connection derivation. Lastly, while experimental validation of derived connections is more limited than in ground-truth simulations, it will be of interest to validate connections derived *in-vivo* using structural connectivity from electron microscopy^26,30^ applied on the same tissue.

## Methods

### Human cortical microcircuit models

We used our previous detailed models of human cortical L5 microcircuits^49^ and L2/3 microcircuits^48^ as ground-truth data where the known connectivity can be used for assessing connectivity derivation performance. Both microcircuit models consisted of 1000 detailed neuron models that reproduced firing rates and patterns in response to step currents^62,79,80^, synaptic properties^6,72,78^, and dendritic sag currents as measured experimentally in human neurons^62,81^. The neurons were radially distributed in a volume with a 250 μm radius and a depth according to human cortical layer dimensions (L2/3 situated 250 μm – 1200 μm below the pia and L5 situated 1600 μm – 2300 μm below the pia)^82^. The models included key neuron types and proportions as measured in humans (L5 circuit: 70% Pyr neurons, 15% SST, 10% PV, and 5% VIP interneurons^71^; L2/3 circuit: 80% Pyr, 5% SST, 7% PV, 8% VIP^71,83,84^), and detailed reconstructed human morphologies. The multi-compartmental neuron models included sodium, potassium, and calcium channel mechanisms in the axon and soma, and h-current in the dendrites, with Hodgkin-Huxley formalism for equations governing activation and inactivation taken from previously published models (see Hay et al., 2011^85^, 2013^86^ for further equations and details). Pyr→Pyr connection probability was 0.15 in L2/3 and 0.09 in L5, within the experimental range from human literature^6,72^. Other connection probabilities were as measured in rodents^78^, and adjusted within experimental ranges to reproduce in-vivo baseline and response firing rates^1,3^ (Table 1). The microcircuit models exhibited emergent power spectral properties of human resting-state EEG which served as an important validation that they captured cortical dynamics^52^.

PV interneurons targeted Pyr basal dendrites, while SST interneurons targeted Pyr apical dendrites. The models used AMPA and NMDA excitatory synapses (τ*_rise,NMDA_* = 2 ms, τ*_decay,NMDA_* = 65 ms, τ*_rise,AMPA_* = 0.3 ms, τ*_decay,AMPA_* = 3 ms), and GABA_A_ inhibitory synapses (τ*_rise,GABA_* = 1 ms, τ*_decay,GABA_* = 10 ms). The synaptic reversal potentials were E*_exc_* = 0 mV and E*_inh_* =-80 mV^87–89^. The microcircuit models received random background excitatory inputs (neuron type specific) using Ornstein-Uhlenbeck (OU) point processes^90,91^. The baseline excitatory OU conductances were set to the following: Pyr = 76 pS; SST = 32 pS; PV = 1,545 pS (due to higher rheobase); VIP = 75 pS in L5, and Pyr = 28 pS; SST = 30 pS; PV = 280 pS; VIP = 66 pS in L2/3. In all microcircuit types and paradigms, the inhibitory component of the OU conductance was set to 0, as the recurrent inhibition in the microcircuit was sufficient to restrain excitability. The data constraints, taken primarily from the middle temporal gyrus and somatosensory cortex, were used to model prototypical cortical microcircuits rather than specific brain areas. All parameters of the L2/3 and L5 microcircuit models involved layer-specific constraints. For more detailed descriptions of human/non-human sources of the model constraints, electrophysiology features optimization, and other parameters such as synaptic conductance and ion channel density values in L2/3 and L5, see Yao et al., 2022^48^ and Guet-McCreight et al., 2023^49^, respectively. The models were simulated using NEURON^92^, LFPy^93^ and parallel computing in high-performance grid (SciNet)^94,95^, using 400 nodes, with a runtime of ∼18 h per 30 minutes of microcircuit simulation. Repositories for simulation code and models of both L2/3^48^ and L5^49^ microcircuits are publicly available.

### Neuronal connectivity derivation

We inferred putative monosynaptic connections by detecting short-lag peaks in cross-correlograms of simulated spike trains (180 minutes in each condition, omitting the initial 1 second of transient activity)^12,13,15,42^. We used previous computational tools (GLMCC, see Kobayashi et al., 2019^42^) for fitting postsynaptic waveform curves to cross-correlograms^42^ for improved derivation. We did a parameter search for the synaptic delay (d_s_), interaction timescale (τ), and flatness of slow fluctuation curve (γ). The synaptic delay determines the start point of the connection function, beyond which the short-lag peak occurs. τ determines the decay timescale of the connection function. The length of τ was chosen that yielded good derivation performance and a good fit of the connection function to the short-lag peak. In the derivation tool, the generalized linear model^42^ is given as:

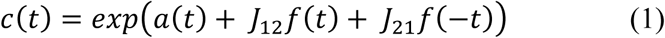

where t is the time from the spikes of the reference neuron, a(t) represents the slow fluctuation curve, and J_12_ and J_21_ represent the connection function in both directions between the two neurons. The time profile of the synaptic interaction is represented as 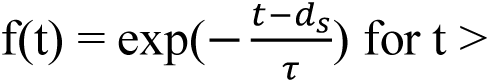 d_s_, and f(t) = 0 otherwise. The confidence interval (J_±_) of J_12_ is:

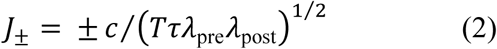

given the observation window (T), firing rates of the pre and postsynaptic neurons (λ_pre_ and λ _post_), and a coefficient c. A connection is considered to exist if significant, i.e. if the estimated connection parameter 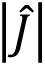 falls outside the confidence interval for the null hypothesis that the connection is absent, given a minimum number of spikes (Tτλ_pre_λ_post_ > 10⁄τ[s^-1^]) in the interaction window (for further details and full set of equations, see Kobayashi et al., 2019^42^). We tested synaptic delays ranging from d_s_ = 1 to 4 ms and slow fluctuation curves ranging from γ = 1 x 10^-4^ ms^-1^ (yielding flatter curves) to 1 x 10^-3^ ms^-1^ (yielding oscillatory curves which fitted the baseline correlations more closely). We used d_s_ = 4 ms and γ = 2 x 10^-4^ ms^-1^ which yielded the best derivation performance and was also the default value set by the tool’s developers based on their multi-parameter optimization methods^42^). We used τ = 4 ms for all connection types, in line with previous studies of cross-correlation analysis^12,13,15^, except SST→Pyr which required a longer τ due to the distal location of the synapses (L5: τ = 8 ms, L2/3: τ = 15 ms). We evaluated derivation performance by calculating the sensitivity, PPV, and specificity^96^:

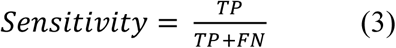

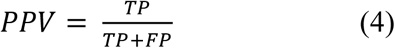

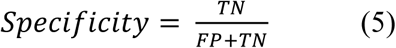

Derivation runtime was ∼30 min per population of 300 neurons, on a standard laptop.

### Silent and low-firing neurons

When deriving connections involving neurons firing < 0.1 Hz as the presynaptic or postsynaptic neuron, the connections were not amenable to the tool analysis due to the lack of spike data, and were thus interpreted as missed connections and given an inferred weight of 0. Silent neurons were defined as firing < 0.2 Hz^97,98^. The proportions of silent neurons in the baseline microcircuit models were 9% in L2/3 and 0% in L5. Low-firing neurons were defined as firing < 1 Hz.

### General neuronal activation

We tested two general activation paradigms of L2/3 Pyr neurons to increase microcircuit activity, by changing background inputs to specific neuron types. In the L2/3 general-Pyr paradigm, background OU conductance in Pyr neurons was increased to 50 pS. In the L2/3 disinhibited paradigm, background OU conductance in SST and PV interneurons was reduced to 28 and 1 pS, respectively, and slightly reduced in Pyr neurons to 24 pS. Other disinhibition paradigms were less useful, e.g. activating VIP interneurons affected mainly SST interneuron inhibition^78,99^ and did not sufficiently reduce the proportion of inactive neurons, reducing SST alone attenuated the overactive population at the expense of an increased proportion of low-firing Pyr neuron population, and reducing PV alone led to a broad distribution with highly correlated firing. A combination of both was thus necessary to achieve the desired mean and distribution.

### Subset neuronal activation

We activated a subset of 100 L2/3 Pyr neurons to induce moderate and jittered spiking, with firing rates similar to that of the L5 microcircuit while minimizing synchronous activity. We adapted our previous models of experimental sensory stimulus responses^48^, whereby each Pyr neuron in the subset was activated using excitatory AMPA/NMDA synapses distributed at 5 locations randomly on the basal dendrites, each with a conductance of 900 pS (corresponding to ∼6 simultaneous presynaptic inputs on top of background input), to yield firing rates similar to L5. The repeated activations occurred in cycles of 50 ms, with jittered stimulus times (post-stimulus delay of 2 – 22 ms), repeating for 180 minutes.

### Correlation tests

Discrete cross-correlations between neuron spike trains were calculated using normalized signals of spike time vectors using 300 seconds of spiking data, with spikes grouped into 3 ms bins. Before correlating, spike time vectors were normalized by subtracting the mean from the signals and dividing by the product of the standard deviation and the vector length to prevent biases due to differences in population firing rates^100–102^.

### Spiking power spectral density

Neuron spike times were converted into binary spike train vectors and then the vectors were summed across all neurons of a given type. Power spectral densities were then computed from the summed spike train vectors using Welch’s method^103,104^ from the SciPy python module with 4s time windows. Power spectral density vectors across random seeds were bootstrapped (n = 500) at each frequency, to compute bootstrapped means and 95% confidence intervals.

### Statistical analysis

We determined statistical significance (*p* < 0.001) using Welch’s t-test when comparing data groups that did not meet assumptions of normality (Omnibus test of normality, *p* < 0.05) and equal variances (Levene test, *p* < 0.05)^105,106^. Otherwise, we used Student’s paired and two-sample t-tests where applicable. When data groups being compared exhibited significantly nonuniform distributions, we additionally used the Mann-Whitney U non-parametric test^107^. We calculated the effect size between two conditions using Cohen’s *d* (dividing the difference of two means by the pooled standard deviation of all values).

## Acknowledgements

We thank the Krembil Foundation for their generous funding support. FM was also supported by the Miner’s Lamp Innovation Fund and by the Department of Physiology at the University of Toronto. MTH was also supported by NSERC and TCAIREM.

